# MethPhaser: methylation-based haplotype phasing of human genomes

**DOI:** 10.1101/2023.05.12.540573

**Authors:** Yilei Fu, Sergey Aganezov, Medhat Mahmoud, John Beaulaurier, Sissel Juul, Todd J. Treangen, Fritz J Sedlazeck

**Affiliations:** Department of Computer Science, Rice University, Houston, TX 77251-1892, USA; Oxford Nanopore Technologies Inc, New York, NY, USA; Human Genome Sequencing Center, Baylor College of Medicine, Houston, TX 77030, USA; Department of Molecular and Human Genetics, Baylor College of Medicine, Houston, Texas, USA

## Abstract

The assignment of variants across haplotypes, phasing, is crucial for predicting the consequences, interaction, and inheritance of mutations and is a key step in improving our understanding of phenotype and disease. However, phasing is limited by read length and stretches of homozygosity along the genome. To overcome this limitation, we designed MethPhaser, the first method that utilizes methylation signals from Oxford Nanopore Technologies to extend SNV-based phasing. Across control samples, we extend the phase length N50 by almost 3-fold while minimally increasing the phasing error by ∼0.02%. Nevertheless, methylation signals have limitations, such as random signals on sex chromosomes or tissue purity. To assess the latter, we also applied MethPhaser on blood samples from 4 patients, still showing improvements over SNV-only phasing. MethPhaser further improves phasing across *HLA* and multiple other medically relevant genes, improving our understanding of how mutations interact across multiple phenotypes. MethPhaser is available at https://github.com/treangenlab/methphaser.

## Main

The emergence of long-read sequencing technologies has enhanced our understanding of the human genome, uncovering novel types of variations between individuals and even tissues^1^. The latest advancements allow us to gain a more comprehensive understanding of single nucleotide variations and more complex structural variations at unprecedented levels. These discoveries reveal novel loci that could potentially impact diseases, evolution, or other important phenotypes^2–6^. Over the past few years, novel computational methods have enabled advancement in all these fields by providing more complete human genomes^7,8^, more comprehensive detection of variants at germline and somatic levels, as well as a more realistic view of the genome by providing phasing information^2,9,10^. Here, phasing refers to the assignment of variants to the two copies of each genome as they are present in human and other mammalian cells^11^. Once this assignment is made, it becomes easier to investigate the consequences of two alleles co-occurring on the same DNA molecule, which can have different impacts on specific genes^12^. We differentiate between cis-relationship, which occurs on the same DNA molecule, and trans-relationship, which occurs on opposite DNA molecules. Identifying the relationship between two or more variants is crucial for many downstream applications.

To perform this phasing, which refers to the relationship between variants, three strategies exist. A population-based phasing strategy leverages the information of co-occurrences of variants across multiple hundreds to thousands of individuals^13^. Thus, while this strategy can easily phase entire chromosome arms, it can only do so on common SNPs in the population, and therefore misses the rare and likely disease-causing variants^14,15^. Another strategy, trio-phasing, involves obtaining parental information and using it to phase a single nucleotide variant (SNV) based on the co-occurrence of the parents^16^. This can also phase rare SNV in the population and produce chromosome-wide phasing but requires additional sequencing of the parents, which is not always available^17^. More importantly, this strategy fails to phase de novo variants, which are often causative in diseases such as intellectual disability^18^ and Mendelian diseases^19^. Lastly, per-read phasing leverages only the linking information of variants that are shared in the same read. While this has the advantage that even a *de novo* variant can be assigned to a specific haplotype, it is highly reliant on the distance between two heterozygous SNVs and the read length. Thus, if the read length is not sufficiently long to span the distance of two heterozygous SNVs, phasing cannot be inferred. While per-read phasing is the most comprehensive method, it heavily relies on the read length and may only obtain regional phase information, which is referred to as a phaseblock. Within a phaseblock region, the phased SNVs are assigned to each haplotype and thus comparable. However, between two phaseblocks, the assignment of SNV to haplotype 1 or 2 cannot be determined as no connection information is available. The aim of this study is to improve phasing to be comprehensive, while obtaining longer phaseblocks and reducing their number.

In recent years, the reduction in cost and increase in yield of long-read sequencing technologies have enabled its use to improve variant detection and to directly improve phasing. Utilizing ever-larger read lengths has resulted in complete phasing as they routinely span the distance of two or more heterozygous SNVs. The average span between two heterozygous SNVs in a human is approximately 1kbp^5^, which long reads easily span. Nonetheless, there are certain regions of the human genome with higher concentrations of homozygous SNV, which can pose a challenge for achieving complete phasing even with long read samples that have an N50 read length of 30 kbp or more.

Another source of information provided by long reads is the methylation signal. Higher or lower methylation is often associated with the inactivation or activation of certain regions of the genome and can be another useful tool for the interpretation of certain genes^20^. Several sequencing methods, like whole genome bisulfite sequencing (WGBS), Hi-C sequencing, and short-read sequencing, have been widely used to analyze methylation signals^21,22^. However, those three methods have several drawbacks too. For instance, bisulfite sequencing is proven to have uneven coverage^23,24^, leading to lower-confidence CpG locations. Short reads sequenced from bisulfite-converted libraries often suffer from poor low alignment scores. On the other hand, long-read technologies have been proven to preserve long-range DNA information without the need for massive pre-processing steps^25–27^. Methylation patterns can vary significantly across different tissues, making them tissue-specific. Recent studies have demonstrated a clear relationship between methylation patterns across haplotypes, with some exceptions. One exception is sex chromosomes, where males have only one copy of the X and Y chromosomes, while females have two copies of the X chromosome (i.e., diploid). The activation and inactivation of one of the X chromosome copies is determined by high methylation of one copy^28,29^. This is done at random and doesn’t follow the haplotype structure. Nevertheless, in theory, this would mean one can leverage methylation signals across stretches of homozygosity for autosomal regions of the genome. By doing so, there is potential to improve upon the variant phasing and be less reliant on the distance between two heterozygous SNVs.

Oxford Nanopore Technologies (ONT) allows individual DNA molecules to pass through a mutated biological nanopore The pores are embedded in a membrane, across which a voltage is applied that generates an ionic current through the pores. The detection of modified bases relies on their unique current signatures, which differ from canonical, unmodified nucleotides, as they pass through the pore^21,27,30,31^. In recent years, many tools have been developed to call 5mC at CpG sites, like Nanopolish^27^, Megalodon^32^, Guppy^33^, DeepMod^34^, and Remora^35^. ONT has combined Remora into its official state-of-the-art basecallers Bonito^36^ and Dorado^37^, and in this study, we are using Remora as our methylation caller. Since the emergence of methylation calling technologies, several methods have been developed for utilizing such information to perform phasing on human genomes, all of which depend on allele-specific methylations. Also, in the same haplotype, base modification probabilities in the same CpG locations are similar. MethHaplo^38^ combines allele-specific DNA methylation and SNVs with bisulfite and Hi-C sequencing data for haplotype region identification. A WGBS-based methylation haplotype block identification method^39^ was also proposed for improving heterogeneous tissue samples and tumor tissue-of-origin mapping from plasma DNA. NanoMethPhase^40^ and PRINCESS^2^ are the other two methods that propose to use SNVs and methylation signals together for methylation phasing. Similarly, ccsmeth^41^ also provides methylation phasing on PacBio circular consensus sequencing data^42^. However, none of the above-mentioned methods provide a path to enlarge phaseblock length and further phase more SNVs on ONT data. They all either perform phasing on distinct data types (short reads and Hi-C reads), or only phase methylation events using phased SNV signals. They do not provide improved SNV phasing or read haplotype tagging results.

This paper investigates the utility of methylation signals for the phasing of SNV and general variations. We show that utilizing methylation can improve the phasing overall and thus be leveraged to assign more reads (i.e., shorter reads) to haplotypes, thereby boosting the ability to call variants. We developed MethPhaser, a tool that operates on a set of already phased variants based on SNVs from, e.g., WhatsHap^11^ or Hapcut2^4^. MethPhaser then utilizes the heterozygous methylation information across the autosomes to connect phaseblocks together and thus improve the overall phasing. We showcase the method on HG002/NA24385, where benchmark data is available based on phased long-read assemblies and variant benchmarks. Here we highlight the performance of MethPhaser genome-wide on autosomal chromosomes, with a special focus on medically relevant genes, especially the *HLA* region^43^, where phasing is most important. To demonstrate the versatility of MethPhaser, we evaluated its performance on various human populations and tissue types. Our results show that MethPhaser improves variant-based phasing with minimal impact on phasing errors. This represents a novel and valuable enhancement for variation analysis.

## Results

### Overview of the MethPhaser benchmarking pipeline

We developed MethPhaser to leverage methylation information to extend SNV-based phasing. This approach is conceptually novel, as we leverage heterozygous methylation signals across stretches of homozygous SNV regions for autosomal chromosomes. MethPhaser takes as input a file containing pre-phased SNVs (e.g. from WhatsHap^11^) along with a BAM file containing methylation tags. MethPhaser is described in detail in the method section. In brief, there are three main steps of MethPhaser (**Figure 1a**): 1. Calculate statistically different methylation CpG locations based on SNV-haplotagged (i.e., labeled) reads. 2. Iteratively haplotag reads with methylation information. 3. Bridge disconnected phaseblocks with newly haplotagged reads. Steps 1 and 2 are repeated until either no more reads in the disconnected regions can be further assigned or the iteration number reaches a user-defined limit (default 10). An illustration of the 3 steps is shown in **Figure 1a**.

**Figure 1.**
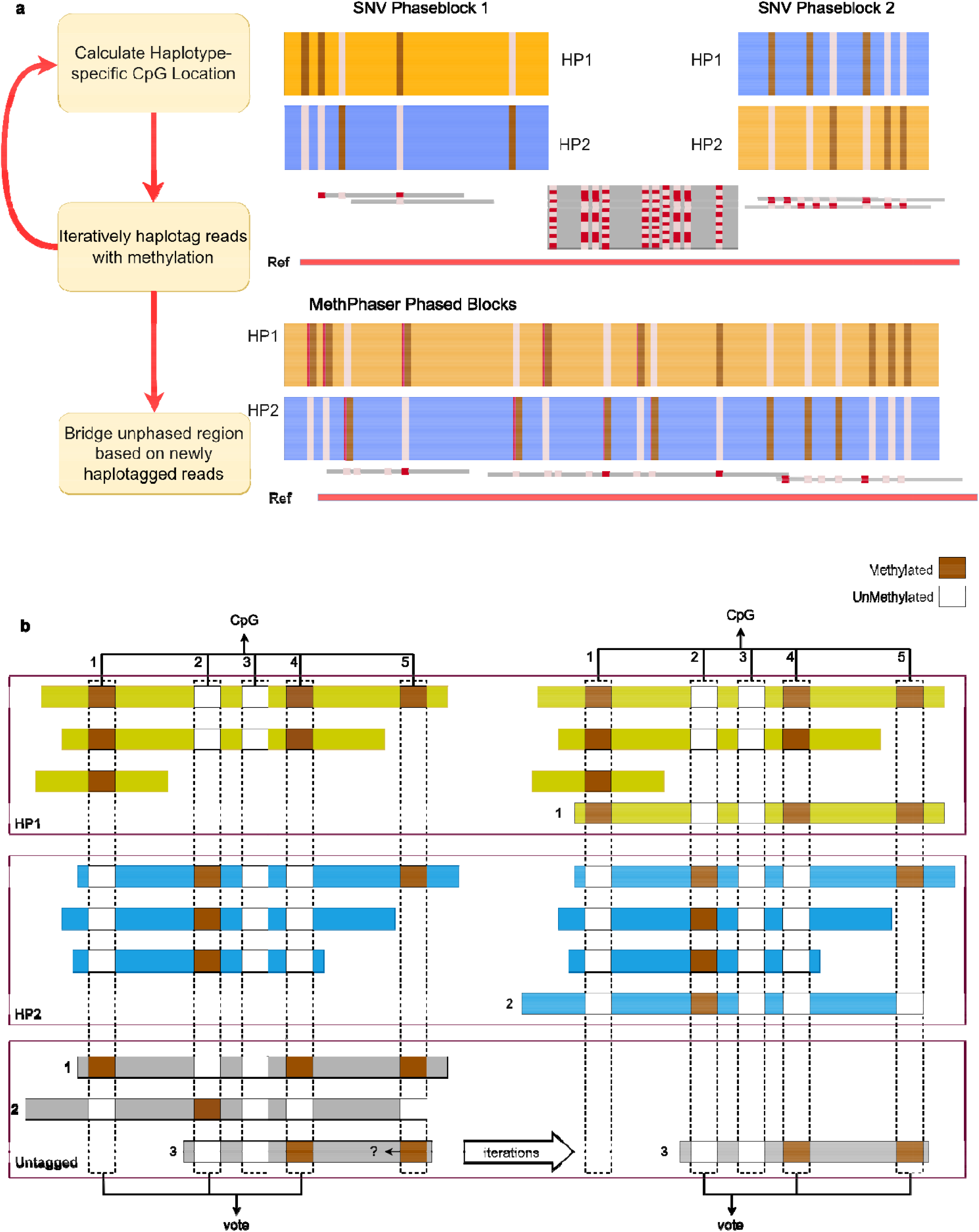
Overview of MethPhaser. **a**, MethPhaser overview. MethPhaser includes 3 major steps: 1. Calculate statistically different methylation CpG locations based on SNV-haplotagged reads. 2. Iteratively haplotag reads with methylation information. 3. Bridge disconnected phaseblocks with newly haplotagged reads **b**, Schematic example of a region where methylation information can help to improve phasing overview. MethPhaser summarizes the methylation patterns in SNV haplotagged reads in each phaseblock and assigns unhaplotagged reads via pattern matching. MethPhaser locates similar patterns on SNV-untagged reads and haplotypes in phaseblocks and assigns matched reads into haplotypes. With newly assigned reads, MethPhaser iteratively updates the existing methylation patterns in the phaseblocks and tags more untagged reads. With more tagged reads, the boundary of the phaseblock can be extended to the unphased region and further close the gap.

In this example, two regions of the genomes are independently phased based on SNV information but couldn’t be connected, either because of a stretch of homozygous SNV or other reasons. Here MethPhaser is able to combine both phaseblocks and thus generate a single larger block by leveraging the heterozygous methylation signal in this region. In addition, MethPhaser tags the unassigned reads to either haplotype to enable a more comprehensive haplotype-specific variant calling. This is particularly important for, e.g., somatic variations, where it is important to understand whether they occur within a haplotype. In the end, MethPhaser produces several outputs: a BAM file with altered haplotype assignment of reads, a vcf file with altered SNV phasing result, and also two files that reveal the phaseblock relationships: CSV files indicate the relationships between SNV phaseblocks, and CSV files indicate previously unhaplotagged reads’ haplotype assignment.

**Figure 1b** shows a schematic example of how MethPhaser works; In **Figure 1b**, five CpG locations (a-e) were identified based on the reference genome and three untagged reads (1-3). We first retrieve the base modification score from the basecaller, and the darker red means the score is high, while the lighter red means the score is low. We use the Wilcoxon rank sum test^44^ to determine which CpG locations have a statistically different score between the two haplotypes. CpG location 3 does not have a statistically different score across the haplotype, so it cannot be counted as a vote. The read assignment is divided into two steps: 1. Read assignment based on SNV phased reads. For the CpG locations a, b, and d, we access the base modification score on untagged reads and see if the score is closer to either haplotype 1 or haplotype 2. In this example, the untagged read 1’s base modification score on location 1 is likely classified into haplotype 1, while untagged read 2 is into haplotype 2. Based on the available votes, untagged reads 1 and 2 can be assigned to haplotypes 1 and 2, respectively. MethPhaser’s default parameter requires a minimum of three votes to determine a read’s haplotype. At least two base modification scores are required for the Wilcoxon rank sum test^44^ (e.g., location e at step 1). Therefore, the untagged read 3 could not be assigned in step 1 because of insufficient votes. 2. After one iteration, the number of base modification scores is sufficient at CpG location e, which makes the vote number of untagged read 3 sufficient. The untagged read 3 can be further assigned to haplotype 1. Finally, MethPhaser goes into the phaseblock assignment step. MethPhaser takes the reads that are assigned by both neighboring extended blocks into votes for the phaseblock relationship assignment. With MethPhaser defined extended boundary, the previously untagged reads can be tagged based on the SNV and methylation information in the first and second extended boundaries. The read is assigned to a switched haplotype in those two boundaries, which indicates the switching relationship between those two neighbor SNV phaseblocks.

### MethPhaser: Methylation as an extension of SNV phasing

To assess the accuracy of MethPhaser, we utilized the Genome in a Bottle (GIAB) SNV benchmark (V4.2.1) and its reported phasing information based on assembly^45^. Here we compared the benchmark phasing results to those obtained using standard SNV-only phasing and the MethPhaser-enhanced phasing using ONT 60X reads from R9 flow cells. We assessed the performance of the SNV and MethPhaser output based on N50 (length of phaseblocks) and phasing errors. We measured these errors as either flip errors (single SNV assigned to the wrong haplotypes) or switched errors (all subsequent SNVs are consistently assigned to the wrong haplotypes) based on WhatsHap compare (see methods for details). MethPhaser was able to extend the N50 phasing information from the standard SNV phasing (see methods) approach by 1.6X times. For the entire HG002 autosomal genome, MethPhaser reduces the number of gaps between the phaseblocks (the continuous region where the relationship between the SNVs is reported) from 3,179 to 1,722 which represents a significant improvement from 1,706,719 N50 to 3,997,227 N50 of phase length. This includes an increase in the flip rate of 0.02% and no switch error increase compared to the SNV phasing alone. The main error mode from MethPhaser is to assign blocks of phasing wrongly, resulting in switch errors or flipping. As we show for HG002, this is a minor 0.02% increase in error rate from 0.03% for SNV-based phasing to 0.05% improved phasing while gaining significantly longer phaseblocks.

We next investigated if MethPhaser requires 60x coverage or can produce similar advancements on different coverage levels. **Figure 2a** shows the N50 comparison based on R9 data across 30, 60, and 80x coverage. Here, 30x represents a single flow cell run. In each case, we could report a significant improvement of SNV phasing based on methylation signals for 30x (1.78 fold), 60x(2.34 fold), and 80x (2.53 fold) coverages. In addition, we also measured the increase in switch error rate, which was from 0% to 0.01% across the different coverage data sets, as shown in **Figure 2b**. Altogether this leads to many more SNV phases, and thus, the number of unphased regions (i.e., gaps) is reduced. **Figure 2c** shows the results, where MethPhaser can reduce the gap number from 81% (R10 30X coverage) to 54% (R9 60X coverage) of the previous SNV-based gap number. Thus, clearly showing an improvement independent of the coverage levels.

**Figure 2.**
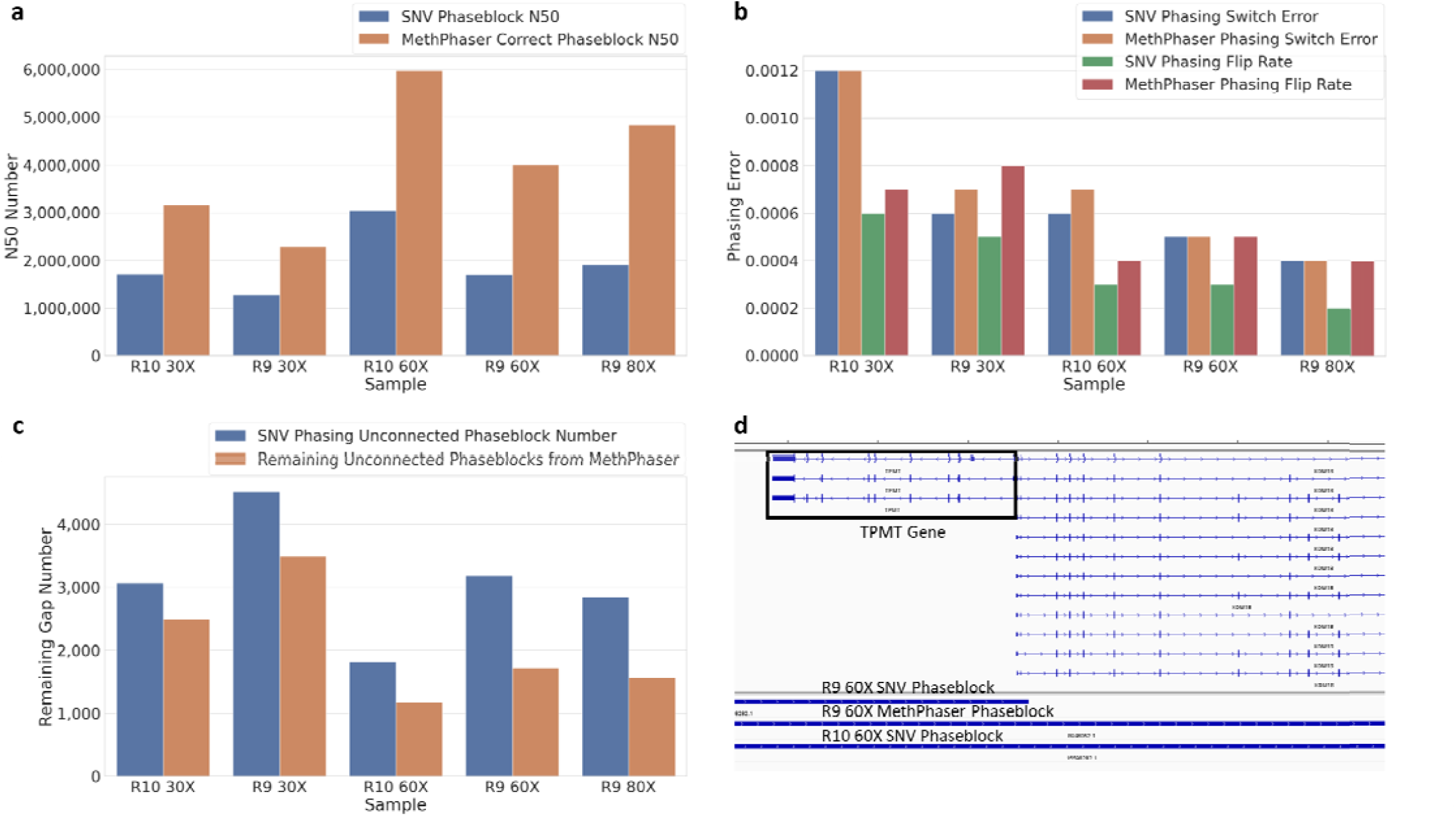
Phasing improvements of MethPhaser across HG002. **a**, SNV Phase N50 and MethPhaser Corrected N50. MethPhaser increases the N50 of phaseblock (falsely connected removed) by 1.6-2.5X. **b**, Comparison of SNV and MethPhaser’s switch error and flip rate. MethPhaser maintains the same level of switch error as the SNV-based method. MethPhaser increases the flip rate by 0.02% over the SNV-based method. **c**, Remaining disconnected gaps on HG002 show that MethPhaser leaves fewer unconnected blocks than SNV phasing methods. With the same coverage, reads from R10 flow cells generally produce fewer gaps. **d**, Improved phasing around the Thiopurine methyltransferase (*TPMT*) gene (associated with encoding the enzyme metabolizing thiopurine drugs) as an important example. An IGV plot showing a *TPMT* example. With R9 60X reads, the SNV-based method cannot fully phase the TPMT region, while MethPhaser is able to phase the *TPMT* gene. The R10 60X SNV-based method can also phase the *TPMT* gene.

Next, we investigated the performance of R10 flow cells from ONT. These represent significant improvements for SNV calling and, thus, potential improvements in SNV phasing itself. We observed that the phasing N50 indeed increases with R10 compared to R9 (see **Figure 2a**, ∼3,000kbp vs ∼2,000kbp). Nevertheless, Methphaser is still able to improve upon SNV-based phasing on R10 data by increasing the N50 by around 1.5 times. Even with only 30X coverage, our program achieves a higher phaseblock N50 (3,152,506) compared to 60x SNV phasing (3,034,229) alone. **Figure 2b** shows the consequences of a small switch error increase (by 0.06%) compared to SNV phasing.

To provide a specific example of why this matters we will now discuss the Thiopurine methyltransferase (*TPMT*) gene. *TPMT* encodes the enzyme metabolizing thiopurine drugs^46^ and includes SNV sites, located 8 kbp apart, that can lead to a reduced functionality of *TPMT*. Given the co-occurrence of SNV on both sites, it is important to determine if they are in cis-relationship, meaning *TPMT* function is reduced, or in transrelationship, which would result in a deactivation of *TPMT*^*47,48*^. The latter has severe implications for a patient and thus could lead to his/her death by administering medical treatment. Given this motivation, we investigated the ability of SNV and methylation-based phasing to obtain the correct results for *TPMT*. We identified differences in the ability of R9 vs. R10 flow cells to phase this important gene entirely. MethPhaser was able to connect the phaseblocks on R9 and thus close the gap, leading to a fully phased *TPMT* gene based on the methylation information. **Figure 2d** shows this across phaseblocks based on the IGV image together with the gene annotation.

Independent of the flow cell and error rates of Nanopore sequencing (R9 vs. R10), MethPhaser shows significant improvements by closing the unphased regions and connecting SNV phaseblocks together (**Figure 2c**), thereby extending phasing across large regions and connecting genes into one continuous phaseblock. When we extend this observation across the entire genome, we indeed see this effect. **Figure 3** shows the N50 increase, switch error, and flip rate of MethPhaser on each chromosome. In **Figure 3a**, we ee that MethPhaser increases phaseblock N50 on each chromosome, except chromosome 15. During the experiment, we discovered there is no SNV phaseblock before the centromere region on chromosome 15, which reduces the N50 increase. **Figure 3b** shows again that the increase results only in a small increase in phasin errors. The largest increase in switch error is 0.0045%, and the largest increase in flip rate is also 0.0045% compared to SNV-based phasing. Thus highlighting that the N50 improvements do, on average, not increase the error rate significantly.

**Figure 3.**
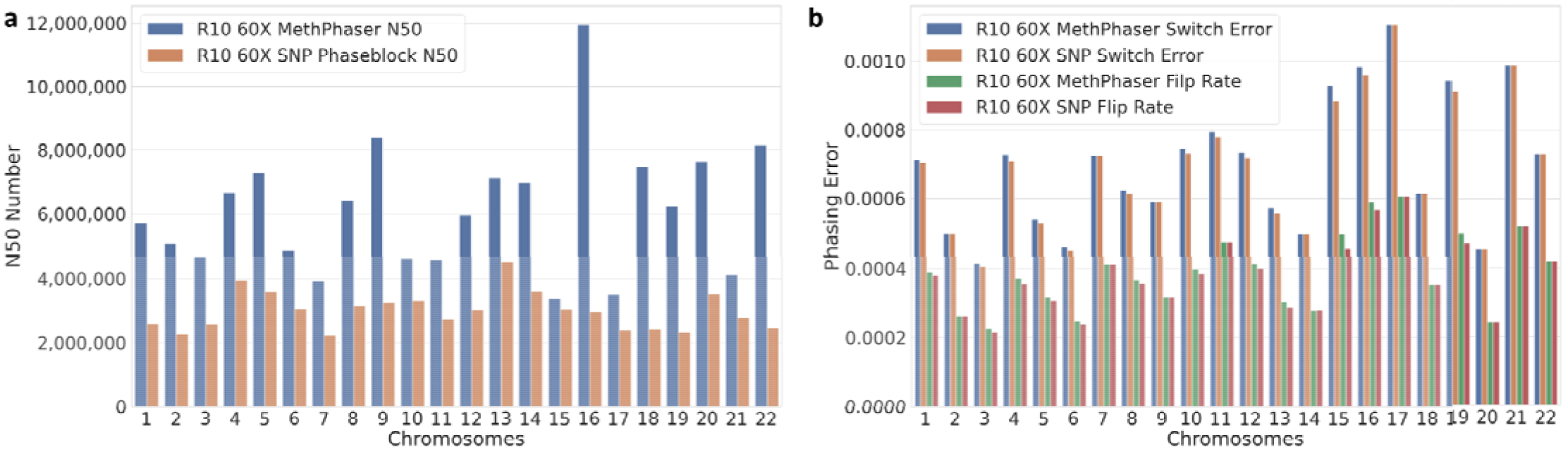
Phasing improvements per chromosome. Per Chromosome N50, the switch error and flip rate of R10 60X reads. **a**, Improvement of phaseblock N50 on each chromosome. **b**, MethPhaser maintains the same switch error and flip rate level.

### MethPhaser improves insights into complex medical genes

Building on our previous genome-wide results and taking into consideration the challenging but medically relevant nature of *TPMT* (thiopurine S-methyltransferase) gene, we sought to evaluate the performance of MethPhaser on this critical genomic region. GIAB has recently published a list and benchmark of 273 medically relevant genes that pose significant challenges to resolve^49^. To determine whether MethPhaser’s improved phaseblock N50 results in a greater number of linked genes compared to SNV-based phasing methods, we conducted a comparative analysis of phaseblock coordinates using MethPhaser, HapCUT2, and the challenging medically relevant genes (CMRGs) benchmark. The comparison result indicates that MethPhaser capable of phasing a greater number of medically relevant genes together, thus allowing for a deeper insight into the counterplay of their respective variants. This is exemplified as we could reduce the phaseblocks across the 273 genes. Overall, MethPhaser could report phasing for 265 (97.1%) of the genes, while SNV phasing was only reported across 258 (94.51%) of these genes. Furthermore, MethPhaser was able to report the phasing with only 140 phaseblocks across the entire set of genes compared to 160 phaseblocks from SNV phasing alone.

To further showcase the importance of MethPhaser and give concrete examples where phasing further matters, we investigated the *HLA* region on chromosome 6. This is a highly complex region of the human genome encoding multiple disease-relevant genes impacting the immune system, diabetes, cancer progression, and many other diseases or general medical phenotypes^50,51^. Thus, a complete phasing helps to interpret the variations across these complex genes of class I and II.

Across the region, MethPhaser was able to extend the phaseblock length by 132,079bp while connecting 3 SNV phaseblocks together. MethPhaser successfully reported larger phaseblocks (validated with GIAB trio-phasing result) that connect both genes. **Figure 4** shows the example of chr6: 30,400,000-31,300,000, which includes the *HLA-E* and *HLA-C* genes in the *HLA* Class I region. We can see in **Figure 4b** that the SNV-based method cannot form a phaseblock that connects those two *HLA* regions, making the phasing status of the *HLA-E* and *HLA-C* unlinked. However, using the methylation information, we are able to link *HLA-E* and *HLA-C* together. The IGV plot shows that previously untagged reads are being tagged by MethPhaser and assigned to haplotype 1 or 2. This also enables improvement in the assessment of variants, as exemplified by a haplotype-specific insertion. The reads with that insertion are mostly clustered into haplotype 1 in this example.

**Figure 4.**
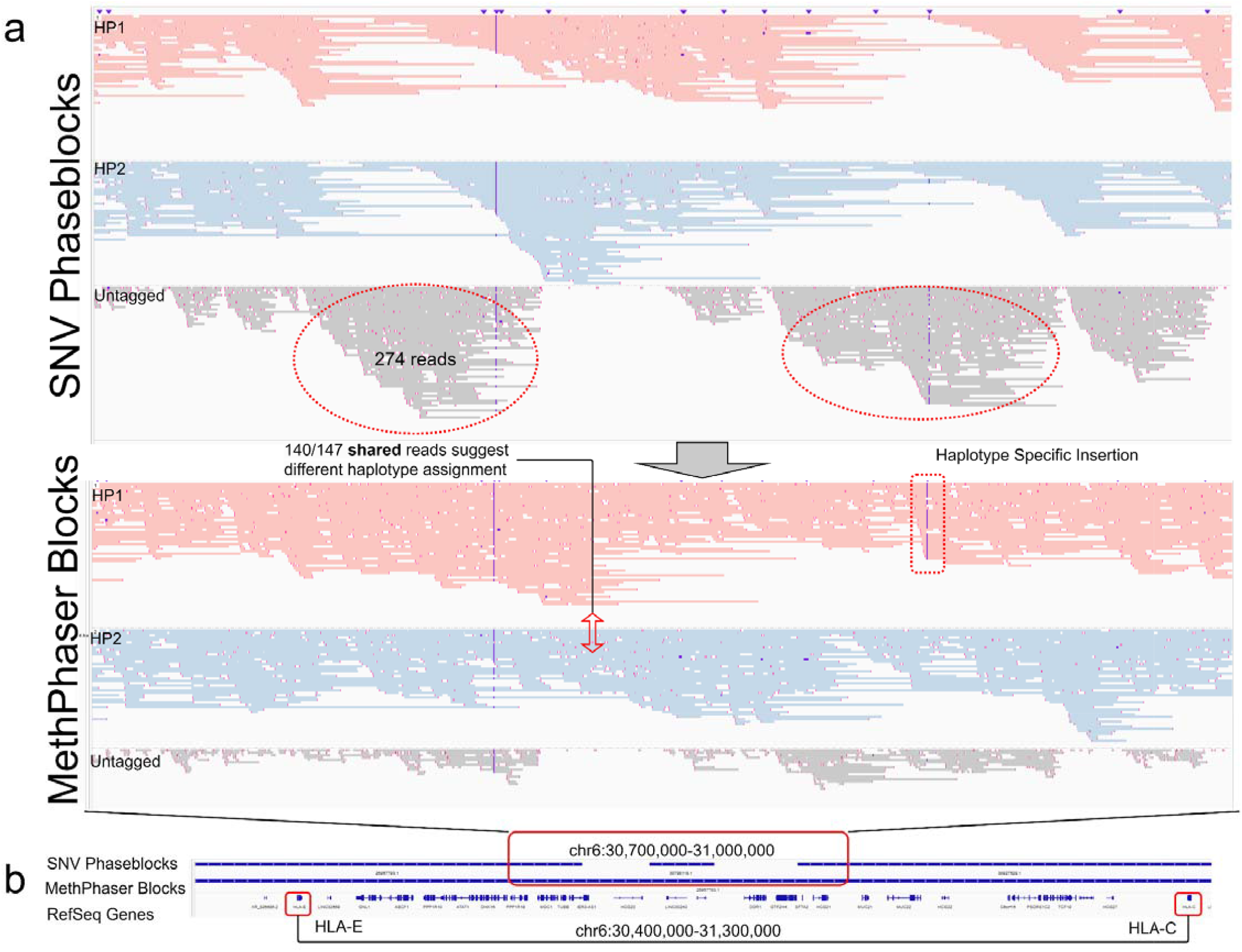
Phasing improvements across *HLA*. **a**, Improved reads assignment connects *HLA-E* and *HLA-C*. **b**, IGV coordinate of the improved phasing regions. The example of MethPhaser improves phasing and read tagging on *the HLA-E* and *HLA-C* genes from the *HLA* Class I region. This example shows chr6 30,400,000-31,300,000, which includes two *HLA* regions. The traditional SNV-based methods cannot form a single phaseblock (with R10 60X ONT reads) that connects two *HLA* regions. However, with MethPhaser, we are able to achieve a single block that covers both regions. With a closer look at the SNV unconnected regions (chr6: 30,700,000-31,000,000), the IGV plot shows that previously untagged reads are tagged by MethPhaser, and MethPhaser discovers a haplotype-specific insertion.

Thus overall, we could highlight the importance of MethPhaser and its improvements for phasing not only genome-wide but also more focused on medically relevant genes and regions (e.g., *HLA*) of the human genome.

### Improved phasing over human population and patient data

In the previous sections, we emphasized the critical importance of phasing and the performance of MethPhaser based on HG002, a cell line where we have benchmark data available. To further validate the robustness and generalizability of MethPhaser, we sought to evaluate its performance across multiple samples from diverse human populations. Furthermore, we wanted to assess the ability to improve phasing even for less optimal ONT data (e.g., shorter reads). To test the effectiveness of MethPhaser across different populations, we investigated its performance across HG01109 (Male, PUR), HG02080 (Female, KHV), and HG03098 (Male, MSL)^52^. We followed the same analysis procedures that were used for HG002 to enable comparability (see Methods). Despite a larger read N50 for these samples (∼40kbp) compared to HG002 (∼30kbp), we observed similar improvements in MethPhaser across all three samples.

Genome-wide SNV-based phasing reported an N50 phase length between 9Mbp and 28Mbp, achieving a higher overall N50 than HG002. This is expected, given the longer read N50 for these samples. Nevertheless, MethPhaser improved upon these N50 phasing lengths in each case, showing an improvement of between 1.69 fold (HG03098) up to 2.55 fold (HG02080) across the samples (**Figure 5a**). This is based connection of several phaseblocks, thereby reducing the number of phaseblocks genome-wide. T on the e most extreme example across the three individuals was HG03098, where MethPhaser reduced the number of phaseblocks from 947 to 643. Furthermore, the inclusion of methylation information reduced the number of phaseblocks in HG02080 from 1,087 to 743. Full details for each sample can be found in the **Supplementary Table**.

**Figure 5.**
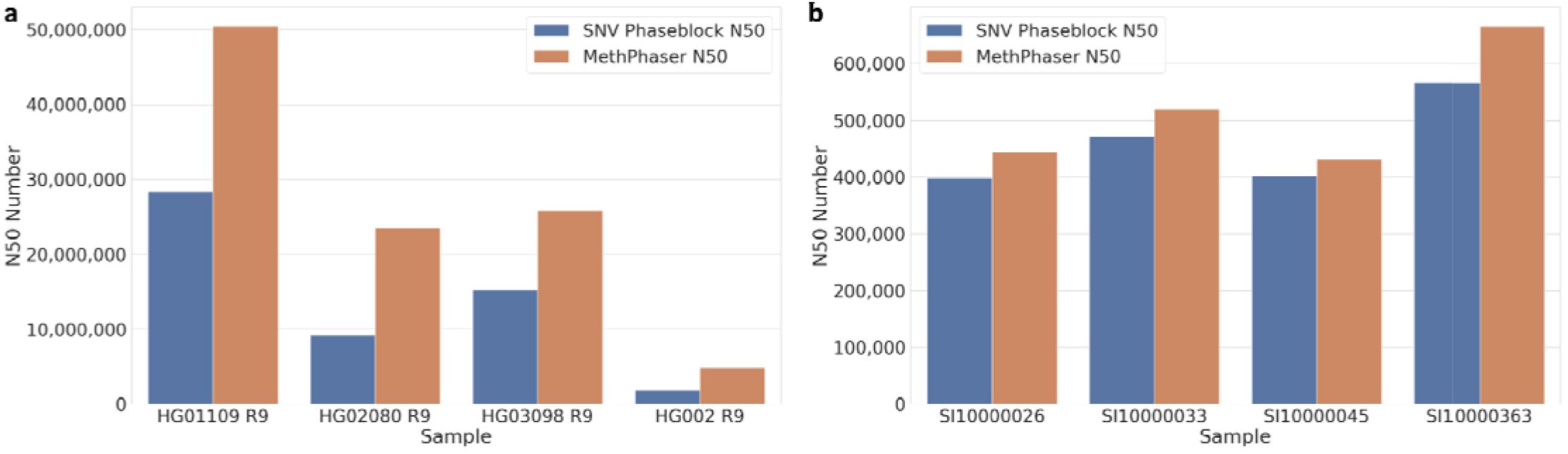
Phasing improvements across different human populations and blood derived samples. N50 increase of HPRC pangenome samples and patient blood samples. **a**, N50 increase in ultra-long, high-coverage pangenome cell-line samples from different ethnic backgrounds. MethPhaser achieved a 1.69-2.55x N50 increase. **b**, N50 increase in patient blood samples from different ethnic backgrounds. MethPhaser achieved a 1.07-1.17x N50 increase due to a much shorter read length.

We have further checked the minimum number of phaseblocks that covers all phaseable medically relevant genes. In HG01109, we could lower the number of phaseblocks to 76 from the initial 82 blocks based on SNV phasing. Two better examples are HG02080 and HG030989, which reduced the phaseblock number across the medical genes from 123 to 113 and 108 to 96, respectively. Given the much longer read length (HG01109 N50: 46,836, HG02080 N50: 41,591), MethPhaser can cover all GIAB-reported complex medically relevant genes on the genome with HG01109 and HG02080 samples, while the SNV-based method failed to do so on HG03098 with the *RHCE* gene on chr1 (MethPhaser can phase) and the *SRR* gene on chr17 (MethPhaser failed to rescue).

Finally, we wanted to assess the performance of MethPhaser on patient samples that might not be as ideal as certain cell lines. For this demonstration, we sequenced two Hispanic and two Caucasian samples with ONT R9 flow cells. Each sample was sequenced with one ONT flow cell, resulting in 24x to 37x coverage. Given the nature of the samples, the N50 is much lower than what was achieved using cell lines (R9 cell line HG002: ∼30 kbp, R9 cell line HPRC: ∼40 kbp, R9 tissue: 12 kbp). Thus, the resulting SNV phasing is overall reduced. Across the four samples, we measured an average N50 of 459.7 kbp based on SNV phasing. This was improved from MethPhaser up to 515.2 kbp average N50 (**Figure 5b**). Given the reduced initial SNV phaseblock size, MethPhaser was still able to improve the overall phasing. The degree of improvement was limited by multiple short SNV phaseblocks that do not allow us to confidently assign reads between methylation and SNV-based phasing. Nevertheless, this demonstrates the ability of MethPhaser to work even on blood samples, where potential tissue heterozygosity makes it harder.

Despite the modest increase in phaseblock N50, we were able to significantly reduce the number of total phaseblocks from 11,916 to 9,837 using MethPhaser. This reduction in the number of phaseblocks has important implications for downstream analysis, as it allows for a more streamlined and accurate identification of disease-relevant variants.

## Discussion

Phasing is an important step for obtaining a more complete picture of genetic variation in the human genome, with about 1-5% of human genes being influenced by these unbalanced DNA sequence variants^12^. In this manuscript, we present a novel method, MethPhaser, that utilizes allele-specific methylation information to improve SNV phasing. This is done by bridging over the gaps introduced by stretches of homozygous SNVs that otherwise cannot be overcome with SNV phasing alone. To showcase this novel approach, we have applied it to multiple cell lines from different human populations and tissue samples including blood-based samples. The method also shows the potential of being applied on other diploid mammals. In each case, MethPhaser was able to not only extend the SNV-based phasing by joining neighboring phaseblocks, but also haplotagged more reads that could not be tagged by SNV-based methods, which has the potential to improve variant calling. By benchmarking against HG002, which has a known truth set, we could demonstrate that this joining of phaseblocks leads to only a minimal increase in phasing error (i.e., switch error). This is due to checks and thresholds used by MethPhaser to uphold an accurate phasing result, rather than just greedily joining phaseblocks together. The incorporation of methylation information comes at no additional costs, as long reads such as Oxford Nanopore Technologies include methylation information without additional preparation or sequencing runs.

The utilization of methylation for phasing is not without limitations, the most obvious of which is the inability to improve phasing for human sex chromosomes. This is because the random deactivation of chromosome X in females would lead to an inconsistent haplotype pattern. Thus, we exclude the X and Y chromosomes from our benchmark. For autosomes, we rely on initial phasing results based on SNVs. Our results, as well as SNV phasing results, are dependent on the accuracy of methylation detection, overall read length and a certain minimum coverage of the data set. This can be easily seen across the cell lines from HPRC, which had a larger read length N50 (∼40 kbp) compared to the blood-derived patient data (12-15 kbp N50). In both cases, MethPhaser could extend and improve the overall phasing, but this was limited in the patient samples by initially smaller phaseblocks. The smaller phaseblocks do not provide enough information to make a clear decision on how they should be phased (e.g. **Figure 1**) and thus MethPhaser does not incorporate them due to the desire to retain a high precision in the phasing accuracy. We further investigated the role of sequencing error or noise in the ability of phasing improvements. Here, we measured the performance of MethPhaser across R9 and R10 flow cells. The latter improves the SNV variant accuracy and thus the phasing overall. Using MethPhaser we could demonstrate the improvement in runs using both R9 and R10 flow cells. Overall, we benchmarked MethPhaser across different conditions, showing that its principles are valid and that it can improve phasing across many medically relevant genes, including the *HLA* region. It is also worth noting that MethPhaser helped the phasing of a haplotype-specific insertion around the *HLA* region in our example (**Figure 4**). SNV-based phasing methods like HapCUT2 and WhatsHap perform poorly around the SV regions^53^, which is likely to be the reason for the phaseblock gap located before the *HLA-C* region. MethPhaser shows the potential of phasing haplotype-specific SVs, but requires more experimental results to show the overall improvement, which could be a future step of the benchmarking.

An interesting point when using methylation signals for phasing is the tissue-specific nature of methylation signals. This makes it hard to predict the performance of MethPhaser across different tissues, as the signals can vary. We have tested MethPhaser across different cell lines and in blood, the results of which suggest that its performance is more driven by read length and sequencing coverage than other factors. Even in the most diverse tissue (blood) that carries cells from other tissue types, MethPhaser improved the phasing results over the SNV-based method. Sequencing a certain tissue type alone (e.g., muscle or skin) should thus only improve the phasing results. In blood, while other tissue types might be present, the overall concentration per tissue is minimal. MethPhaser relies on a majority vote to make decisions about combining existing phaseblocks, which is often rather conservative but leads to fewer errors as we could show in the benchmark (**Supplementary Table**). In future work, we plan to leverage and improve the detection of heterogeneous signals in the methylation data to improve phasing also for cancer genomes where larger amplifications of regions will carry different methylation signals. Here, MethPhaser could further significantly improve our phasing and thus understanding of cancer evolution.

Overall, MethPhaser is a novel approach to utilize 5mC methylation signals to improve phasing and thus delivers more key insights into the co-occurrence of mutations across medically important genes.

## Methods

### MethPhaser: phasing based on methylation signal

The primary input for MethPhaser consists of a VCF file containing phased SNVs and a tagged BAM file containing methylation information. To perform an extended phasing based on SNVs, three main steps of MethPhaser are taken: 1. Identify haplotype-specific methylation in each SNV phaseblock with SNV haplotagged reads. 2. Iteratively assign unphased reads in extended SNV phaseblocks based on methylation signals in the unphased reads. 3. Infer the relationship between neighboring extended SNV phaseblocks with methylation haplotagged reads. The CpG probability reaches beyond the phaseblocks, and to make sure that we assign enough reads, we expand our target boundary from the end of the last SNV phaseblock to the start of the next SNV phaseblock. We define SNV phaseblock as *B={b*_*1*_, *b*_*2*_, *… b*_*n*_*}*, and the start and end of the SNV phaseblock is *S={s*_*1*_, *s*_*2*_, *…, si, …, s*_*n*_*}* and *E={e*_*1*_, *e*_*2*_, *…, e*_*n*_*}*. The extended phaseblock, *Be={be*_*1*_, *be*_*2*_, *…, be*_*i*_, *…*, *be*_*n*_*}*, start and end of extended phaseblock will then be *Se = {s*_*1*_, *e*_*1*_, *…, e*_*i-1*_, *… e*_*n-1*_*}, Ee = {s*_*2*_, *s*_*3*_, *…, s*_*i+1*_, *…, s*_*n*_, *e*_*n*_*}*.

### Infer haplotype-specific methylation in extended SNV phaseblock with SNV haplotagged reads

Our method begins with determining whether each CpG location in an extended SNV phaseblock is haplotype-specific. This involves collecting base modification scores from SNV haplotype-tagged reads and calculating base modification probabilities for each CpG location, as shown in **Figure 1b**. The probabilities are the number of MM tags reported from Remora called BAM files. We use the Wilcoxon rank sum test^44,54^ to determine whether the two allele’s base modification probabilities are statistically different. Subsequently, MethPhaser generates a list of CpG locations with statistically different base modifications between haplotypes in the same allele, which are considered allele-specific methylations. Due to the fact that those allele-specific methylations are located in the same extended SNV phaseblock, those are naturally considered haplotype-specific methylations.

### Iteratively assign untagged reads in extended regions

We further assign the reads that cannot be phased by the SNV-based method by collecting their base modification probabilities in pre-calculated allele-specific CpG sites in step 1 and to determine if the probabilities are closer to either haplotype. To ensure better accuracy, we set a minimum coverage of each haplotype (default 3), and a minimum number of votes (default 3) are required for a read’s haplotype assignment. For instance, in **Figure 1b**, we can see in the unphased read 1 that the first CpG site has a similar probability to the haplotype 1, also the second CpG site. Those CpG locations are treated as votes, and if more votes within one read support it as either haplotype (above threshold), we will assign the read as that haplotype. We disregard the third CpG site on unphased read 1, as it has minimal difference between the two haplotypes, and the last CpG site, which lacks sufficient coverage of haplotypes 1 and 2. Based on the three remaining votes, they all suggest haplotype 1, so MethPhaser assigns the unphased read 1 to haplotype 1. Similarly, we assign the unphased read 2 to haplotype 2.

After the read assigning with SNV-phased reads, MethPhaser assigns unphased reads in unphased regions iteratively. In the previous example (**Figure 1b**), after MethPhaser assigned unphased reads 1 and 2 into haplotype 1 and 2, we can see that the unphased read 3’s haplotype was previously unable to decide since there was not enough information on its last CpG location. However, the coverage of haplotypes is sufficient due to our newly haplotagged reads (**Figure 1b**, first iteration), and thus, un-haplotagged read 3 can be assigned. Moving on, we update the list of haplotype-specific methylation and haplotagged reads to assign unphased reads further until no more reads can be assigned or the iteration number reaches a user-defined number.

### Infer neighbor SNV phaseblocks’ relationship from meth-phased reads

Finally, with all these newly haplotagged reads, we can infer neighbor SNV phaseblocks’ relation from our MethPhaser haplotagged reads. No haplotype-specific SNV information in the unphased regions leads to consistent haplotype assignments in neighboring regions, as shown in the example in the **supplementary Figure 2**. However, if those previously un-haplotagged reads are haplotagged by both of the two neighboring extended phaseblocks with the SNV and methylation information, they can bridge those SNV-based phaseblocks together. The task is to look for reads assigned to haplotypes via methylation data across inbound extensions of consecutive regions. If the haplotype assignment of the same read is switching, MethPhaser will record a vote of switching relationship between two neighboring phaseblocks. By default, MethPhaser ignores the largest gap between the SNV-based phaseblocks, which is the centromere region, to save computation time.

### Post-processing and result filtering

To reduce the false positive rate, we also applied a filtering script that allows users to determine the minimum reads and voting confidence (the difference between the reads supports “same” and “not same”) that support the relationship calls. We provide two parameter suggestions based on the experimental results: 1. Best success rate of connecting SNV phaseblocks, with no limitation of minimum reads and voting confidence; 2. Best accuracy, with *read coverage/(genome-wide block number/1000)* as a minimum read number supporting block relationship assignment requirement and more significant than *0*.*5* voting confidence. MethPhaser default uses the best accuracy parameter. The details of the benchmarking process are described in the benchmarking section.

### Output files

The output files contain multiple CSV files that indicate the neighbor relationships and previously un-haplotagged read assignments. Based on those two CSV files, we further modify the BAM and VCF files with pysam^55^ and samtools^56^ for the user.

1. **BAM file**. The generation of the BAM file depends on the two CSV files mentioned above. Given the neighbor relationship CSV files, MethPhaser generates a list of extended blocks that need to be flipped, i.e., switch haplotype 1 or 2 assignment. Three types of reads need to be processed: **a**. The reads that tagged by SNV-based methods. Those reads are flipped if they are in that list; otherwise, they will be output without modification. **b**. The reads that were tagged by MethPhaser. Those reads are output with their new haplotype assignment, and if the block is in the block-flipping list, the haplotype assignment is switched during the output. **c**. the reads tagged by MethPhaser but overlapped with the previous block. Those reads are removed in this block’s read assignment process since they’ve already been output with the previous block. The reads assignments are from the read assignment CSV files.
2. **VCF file**. Similar to the BAM file, the VCF files are generated by considering the neighbor relationship CSV files. MethPhaser reuses the block-flipping list to determine which block’s SNVs’ haplotype assignments need to be switched.
3. **Neighbor relationship CSV files**. Neighbor relationships are stored in CSV files, which are separated by chromosomes. In the CSV file, the header indicates the SNV/extended phaseblock, its next SNV /extended phaseblock, and their relationship. The program outputs the true relationship if the truth-phased VCF file is given. Also, the program outputs the read number that supports such relationship assignments.
4. **Previous un-haplotagged read assignment CSV files**. This file is separated by each extended phaseblock. Each CSV file indicates the previous un-haplotagged reads from SNV-based methods’ new haplotype assignment.

### Benchmarking of phasing performance

The benchmarking process includes calculating phaseblock N50, examining the correctness of neighbor SNV phaseblock connection, and calculating the haplotagged reads’ number. **Figure S1** shows an overview of the MethPhaser benchmarking pipeline.

The phaseblock N50 is calculated as the minimum phaseblock length, where the sum of its phaseblocks with all larger phaseblocks spans ≥50% of the total phase length^57^.To make sure the phaseblock connection is as accurate as possible, we chose our “high accuracy” parameter setting for block connecting. On top of that, we excluded the regions we did not connect correctly to get the correctly connecting N50 with HG002 samples based on the comparison of the SNV-based phasing method and the trio-phased VCF provided by GIAB. We compared the correctly connecting N50 with the final phaseblocks produced by HapCUT2.

The haplotagged reads are calculated with SAMtools, which provides the ability to count the number of reads with certain tags. In our study, haplotagged reads are tagged with the “HP” tag, and we compared the number of reads with that tag before and after MethPhaser. Furthermore, to show MethPhaser can haplotag reads in homozygous regions, we also visualized those regions in IGV to have an intuitive result (**Figure 3**).

The entire phasing process starts from the input of raw reads from ONT. The state-of-the-art basecaller, Bonito^36^, Guppy^33^, or Dorado^37^ from ONT, can basecall raw ONT reads with methylation information included. We then used minimap2 v2.24^58^ to align reads to the reference and keep the MM and ML tags. Clair3 v0.1-r12 ^59^ was used for variant calling with the consideration of different read types and fed the variants into HapCUT2^4^ for SNV phasing. We also filtered the allele frequency (AF>0.25) and limited our interested region to “high confidence” regions provided by GIAB to remove potential Clair3’s false positive calls. The WhatsHap *stats* module was used to visualize and generate readable GTF files containing SNV phaseblocks. The WhatsHap *haplotag* module was applied to tag reads based on HapCUT2’s SNV phasing results. MethPhaser takes WhatsHap haplotagged reads, HapCUT2 phased SNVs as input, and performs methylation-based read phasing and SNV-phaseblock chaining. To check the accuracy of our phaseblock connection, we need to determine the true relationship between two neighboring SNV phaseblocks. Two neighboring SNV phaseblocks are considered to have the same haplotype assignment if their haplotype assignments’ relationship against the truth VCF is the same. For example, if haplotype 1 in SNV phaseblock 1 is haplotype 2 in the truth VCF file while the SNV phaseblock 2 has the same relationship, i.e., the haplotype 1 in SNV phaseblock 2 is also haplotype 2 in the truth VCF, we consider the SNV phaseblock 1 and SNV phaseblock 2 are having the same haplotype assignments. Vice versa, two neighboring SNV phaseblocks’ have switched relationships means their haplotype assignment relationships to the truth VCF are also switched. Given these truth relationship assignments, we can easily know the correctness of MethPhaser’s each SNV phaseblock connection with our relationship output and further get the correct N50 on the HG002 sample in **Figure 2**. WhatsHap *compare* was used on checking the flip error and switch error on HG002. The detailed parameters are listed in the supplementary table.

### Statistics and reproducibility

All analyses were performed with the same parameters as the benchmarking section stated.

## Data Availability

The HG002 kit 10/R9 dataset is available at https://labs.epi2me.io/gm24385-5mc-remora/, called by Bonito base caller with profile dna_r9.4.1_e8_sup@v3.3. The reference genome is hg38 from Genome in a Bottle (GIAB NIST). The reads are at the coverage of 80x, and to test the effectiveness of our method in lower coverages, we also randomly subsampled the reads into 60x and 30x.

The HG002 kit 14/R10 dataset is available at https://humanpangenome.org/data.html. It is sequenced with Oxford Nanopore kit 14 (400bps speed) and pore version r10.4.1 and basecalled with Dorado v4.0.0 SUP model + Reomra. The reference genome is also hg38. The reads are at the coverage of 60X, and we also subsampled it into 30X.

Pangenome datasets’ raw reads are available at https://github.com/human-pangenomics/hpgp-data. The raw reads are re-basecalled with Dorado + Remora the R9.4.1 data (SUP model, 5mCG modifications). The reference genome is also hg38. The samples’ coverages are various, around 60X.

## Code Availability

MethPhaser was written in Python3. The software MethPhaser V0.0.1 that was used in this paper and the script that was used to generate results are available online through GitHub at https://github.com/treangenlab/methphaser under the MIT License.

## Acknowledgments

F.J. was part supported by NIH: UM1HG008898, 1U01HG011758-01, T.T. was supported in part by Centers for Disease Control (CDC) contract 75D30121C11180, NIH NIAID P01AI152999. Y.F. was supported in part by funds from Rice University and Ken Kennedy Institute Computer Science Engineering Enhancement Fellowship, funded by the Rice Oil Gas HPC Conference.

## Author contributions

Y.F., S.A. and J.B. developed the initial concept; Y.F. developed the software and performed the experiments and analyzed the data from S.A. and F.S.. M.M. performed the experiment on the blood sample. F.S., T.T., and Y.F. wrote the manuscript. J.B. and S.J. proofread the manuscript. All authors discussed the results and approved the manuscript.

