## Supplementary Figures for "MethPhaser: methylation-based haplotype phasing of human genomes"

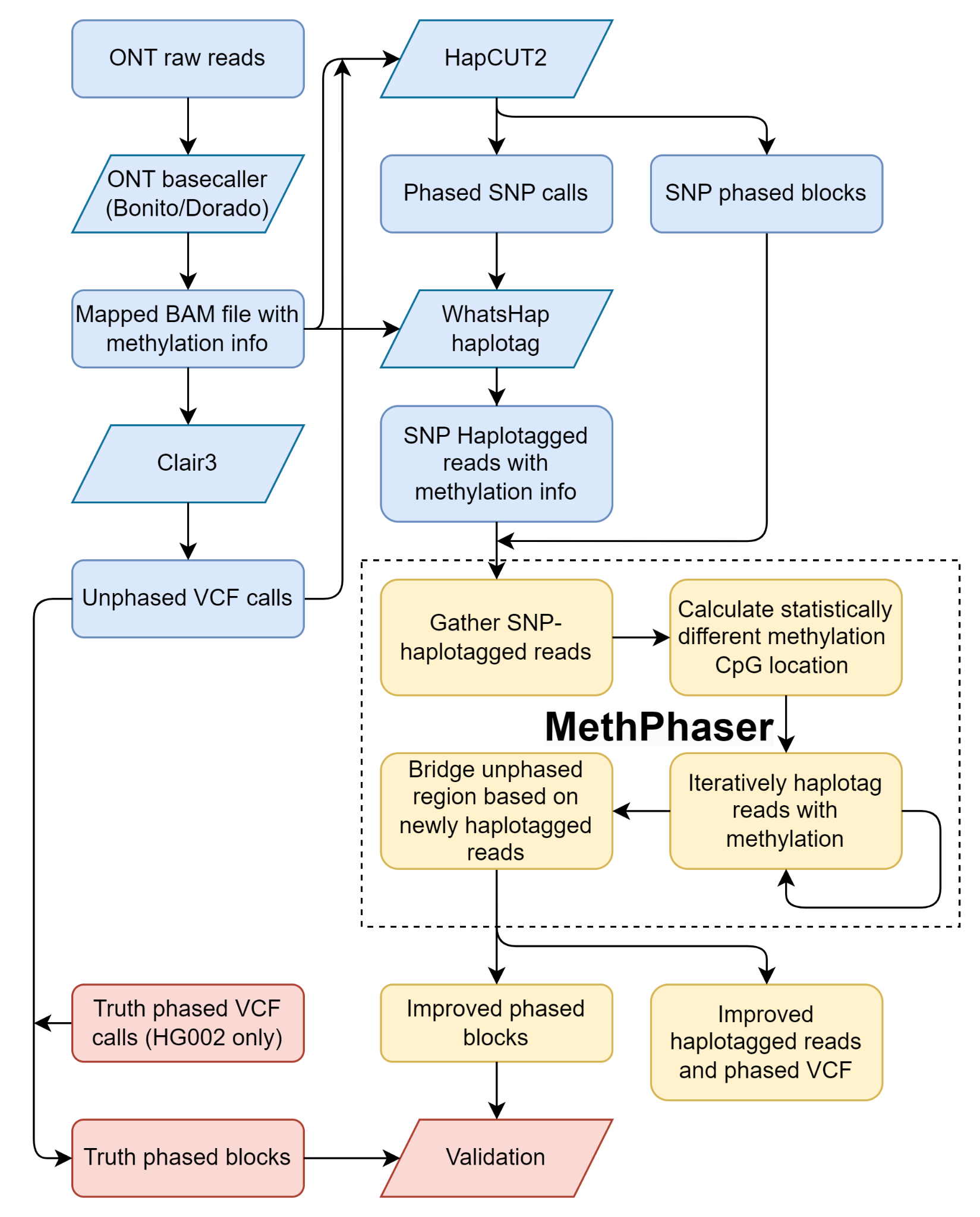

**Figure S1**. Flowchart of MethPhaser Benchmarking Process. Blue blocks represent the traditional SNV-based phasing process, which is the SNV phasing that was used in this work. The yellow blocks, MethPhaser, were directly attached to the traditional SNV-based phasing pipeline for improvement. Red blocks represent the validation process we took in this paper with HG002;

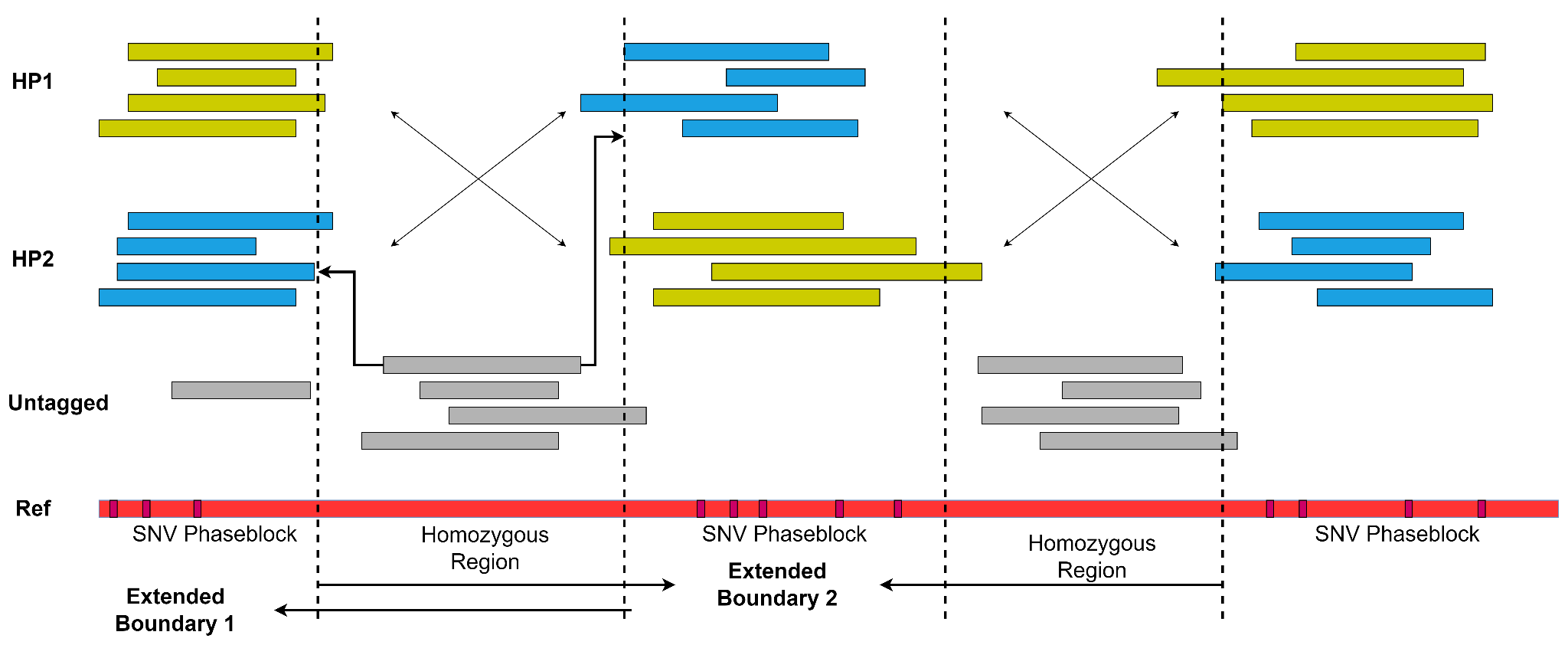

**Figure S2:** The schematic illustration of MethPhaser’s block connection process. We defined several extended boundaries for our classifier for each phaseblock (see method). And we now can start to infer the relationship between the neighbor phaseblocks. Each phaseblock has its own assignments for untagged reads, and the reads in unphased gaps can be assigned by both classified built from phaseblocks based on our definition of extended boundary. So for each previous untagged read that can be tagged by the classifiers we built from the previous SNV phaseblock and the next SNV phaseblock, we can check if those classifiers are outputting the same haplotype assignment or opposite haplotype assignment. If multiple reads support the opposite haplotype assignment, we can infer that the two SNV phaseblocks are having switched haplotype assignment relationship, and further close the gap. Or if the reads support the same haplotype assignment, we can infer that the two SNV phaseblocks are having the same haplotype assignment.
